# The influence of ceftriaxone, ceftazidime-avibactam, and piperacillin-tazobactam on the gut microbiota

**DOI:** 10.1101/2025.04.19.649511

**Authors:** Guohang Huang, Lili Cai, Jinlin Guo, Ruyi Zhu, Lina Zhu

## Abstract

**Purpose:** The administration of antibiotics can induce dysbiosis of the gut microbiota, leading to diseases, e.g., irritable bowel syndrome, metabolic syndrome, and *Clostridium difficile* infection. However, the specific effects of different β-lactam antibiotics on the dysbiosis of the gut microbiota remain poorly understood, particularly in human studies.

**Methods:** This study assessed the impacts of ceftriaxone, ceftazidime-avibactam, and piperacillin-tazobactam on the diversity, composition, and bacterial interactions within the gut microbiota at days 1, 6, and 37 after administration.

**Results:** All three antibiotics significantly altered beta diversity of the gut microbiota by day 6, with ceftriaxone showing the most prolonged effects. Changes in the composition of the gut microbiota were more similar between the ceftazidime-avibactam and piperacillin-tazobactam groups and differed markedly from those in the ceftriaxone group. Consistent with beta diversity changes, bacterial interaction networks showed greater and longer-lasting disruptions of bacterial interaction in the gut microbiota in the ceftriaxone group compared to the ceftazidime-avibactam and piperacillin-tazobactam groups.

**Conclusion:** These findings highlight distinct patterns of microbiota disruption following ceftriaxone, ceftazidime-avibactam, and piperacillin-tazobactam treatments and provide insights for mitigating dysbiosis of the gut microbiota during β-lactam therapy.

## Introduction

The gut microbiota plays a significant role in host nutrient metabolism, immune regulation, and the maintenance of barrier functions [1]. However, while the widespread use of antibiotics has been effective in treating bacterial infections, it has inevitably disrupted the balance of the gut microbiota [2]. Antibiotics inhibit or eliminate susceptible bacteria, leading to reduced microbial diversity and alterations in community composition particularly in the gut microbiota [3], thereby disrupting the homeostasis of the microbiota. Such disturbances may trigger various health issues, including but not limited to irritable bowel syndrome, metabolic syndrome, *Clostridium difficile* infection, and an increased risk of antibiotic-resistant strains [4].

β-lactams are a widely used class of antibiotics that inhibit bacterial cell wall synthesis, making them effective against a broad spectrum of bacterial infections. Among β-lactams, ceftriaxone is a third-generation cephalosporin with a long half-life (5.8 to 8.7 hours) [5,6]. It is effective against a wide range of Gram-positive and Gram-negative aerobic bacteria. Ceftazidime-avibactam combines a third-generation cephalosporin with a β-lactamase inhibitor, expanding its activity against multidrug-resistant Gram-negative pathogens [7]. It is particularly useful for treating complicated intra-abdominal infections, urinary tract infections, and hospital-acquired pneumonia. Piperacillin-tazobactam is a penicillin/β-lactamase inhibitor combination with broad-spectrum activity against both Gram-positive and Gram-negative bacteria, including *Pseudomonas aeruginosa* [8]. It is frequently employed in treating severe infections, such as intra-abdominal infections, febrile neutropenia, and bloodstream infections. These β-lactams remain critical in managing diverse infections and combating antimicrobial resistance. It’s worth noting that the half-lifes of ceftazidime-avibactam (∼2 hours) [9] and piperacillin-tazobactam (1-2 hours) [10] are shorter than ceftriaxone.

The impact of antibiotics on the gut microbiota is theoretically influenced by their mechanisms of action and spectrum[4]. However, most previous studies focus on the antibiotic spectrum against specific pathogens rather than examining their effects on the entire human microbiota or investigate microbiota using animal models. In human microbiota research, two studies has demonstrated that treatment with ceftazidime-avibactam and piperacillin-tazobactam results in lower alpha diversity in the gut microbiota compared to ceftriaxone, implying that ceftazidime-avibactam and piperacillin-tazobactam may inhibit a broader range of bacterial species, thus having a wider antibiotic spectrum than ceftriaxone [11,12]. Additionally, the use of ceftazidime-avibactam has been shown to reduce the relative abundance of *Escherichia coli* and other *Enterobacteriaceae* while increasing enterococci [13]. Similarly, treatment with piperacillin-tazobactam inhibits several species, including *E. coli*, and genera such as *Bacteroides, Bifidobacterium*, and *Veillonella* [14]. However, the comparative effects of these β-lactams on the composition of the gut microbiota, particularly in a longitudinal cohort, are still not well understood.

Following the cessation of antibiotic use, the gut microbiota undergoes a recovery process [15]. However, this process may vary depending on the type of antibiotic treatment administered. One study shows that the gut microbiota are recovered after 1.5 months of treatment by meropenem, gentamicin and vancomycin [16]. The recovery of the gut microbiota from dysbiosis induced by ceftriaxone, ceftazidime-avibactam, and piperacillin-tazobactam remains obscure in the human body.

This study analyzes the effects of ceftriaxone, ceftazidime-avibactam, and piperacillin-tazobactam on the gut microbiota on days 1, 6, and 37 after administration, focusing on comparing their impacts on microbial diversity, compositional changes, and bacterial interaction networks. The recovery of the gut microbiota after treatment by these antibiotics are investigated. These findings will enhance the understanding of dysbiosis of the gut microbiota induced by ceftriaxone, ceftazidime-avibactam, and piperacillin-tazobactam and can be potentially beneficial for providing evidence for rational antibiotic use with less side effects caused by microbiota changes.

## Materials and methods

### Data preprocessing

The raw 16S rRNA sequencing data was obtained from the NCBI SRA database (https://www.ncbi.nlm.nih.gov/sra) under the BioProject “PRJNA922086”. Quality control and sequence trimming, paired-end read merging, and removal of human reads were conducted using the fasterq-dump, flash, and Bowtie 2 softwares, respectively. Pre-treated reads were aligned to the Greengenes2 database (https://greengenes2.ucsd.edu/) for taxonomic classification [17]. Taxa with a relative abundance of at least 0.1% (or 0.01%) in a minimum of 5% (or 15%) of the samples were retained in the final 16S rRNA feature table. Samples with fewer than 1,200 total reads were excluded based on alpha rarefaction results (data not shown).

### Beta diversity analysis

The 16S rRNA feature table was rarefied to match the sequencing depth of the sample with the fewest reads (≥1,200). Beta diversity was computed using the Bray-Curtis dissimilarity metric in the R package vegan [18] and visualized using the ‘prcomp’ function in R. Differences in beta diversity were statistically tested using the PERMANOVA-based ‘adonis2’ function in vegan.

### Differential abundance analysis

Differential abundance of taxa was assessed using LefSe through the ‘run_lefse’ function in the microbiomeMarker package in R.

### Bacterial interaction network analysis

Spearman’s correlation was used to evaluate associations between taxa pairs. The resulting *P*-values were corrected for multiple comparisons using the Benjamini-Hochberg method. Significant correlations (FDR ≤ 0.05) were used to construct unweighted bacterial interaction networks in Gephi [19]. The degree of each bacterial node was quantified using the ‘Avg. Path Length’ function, while modules within the networks were identified using the ‘Modularity’ function. Nodes were color-coded according to their assigned module.

### Institutional review board statement

The study received approval from the Ethics Committee of the Third People’s Hospital of Cangnan County (No. II20250001). The publicly accessible dataset was sourced from the NCBI database (refer to Data availability for details), and information about informed consent to participate is available in the original publication linked to these dataset.

## Results

### Study Cohort

The data for this study were derived from the research conducted by Messaoudene et al. [20], which primarily focused on the ability of a colon-targeted adsorbent, DAV132, to mitigate the effects of three β-lactam antibiotics, i.e., ceftriaxone, ceftazidime-avibactam, and piperacillin-tazobactam, on the gut microbiota. However, the differential impacts of these three antibiotics on the gut microbiota, particularly across different time points, have not been investigated. Therefore, this study focused on the effects of these three antibiotics on the gut microbiota, with an emphasis on comparing the recovery of the gut microbiota after treatment with each antibiotic.

Samples were collected on day 1 (the same day as antibiotic treatment), day 6 (six days after treatment), and day 37 (37 days after antibiotic administration), as these time points had the highest and most comparable numbers of samples (59, 53, and 64 samples, respectively), while other time points had significantly fewer samples (all < 20) (Supplementary Dataset 1). To ensure statistical power was not compromised by sample size disparities, only samples from days 1, 6, and 37 were included in this analysis.

### Effects of ceftriaxone, ceftazidime-avibactam, and piperacillin-tazobactam on beta diversity of the gut microbiota

The impact of antibiotics on beta diversity of the gut microbiota was visualized using PCoA plots and assessed for statistical significance using the Adonis test [18], a type of the PERMANOVA analysis. Two Adonis models were employed: one evaluated the overall effects of the four groups (control and three antibiotics), while the other performed pairwise comparisons between groups.

On day 1, no significant differences in beta diversity were observed among the four groups, indicating comparable baseline levels (Figure 1a). By day 6, beta diversity differences were most pronounced, with the overall Adonis test yielding a *P*-value of 0.002 (Figure 1b). Pairwise comparisons revealed significant differences between each antibiotic-treated group and the control group (P ≤ 0.05), indicating that all three antibiotics significantly altered gut microbiota beta diversity. Moreover, the differences between the ceftriaxone group and both the ceftazidime-avibactam and piperacillin-tazobactam groups were more significant (P ≤ 0.01) than the differences between the latter two groups (P ≤ 0.05). This suggests that on day 6, the gut microbiota profiles of the ceftazidime-avibactam and piperacillin-tazobactam groups were more similar to each other than to the ceftriaxone group.

**Figure 1.**
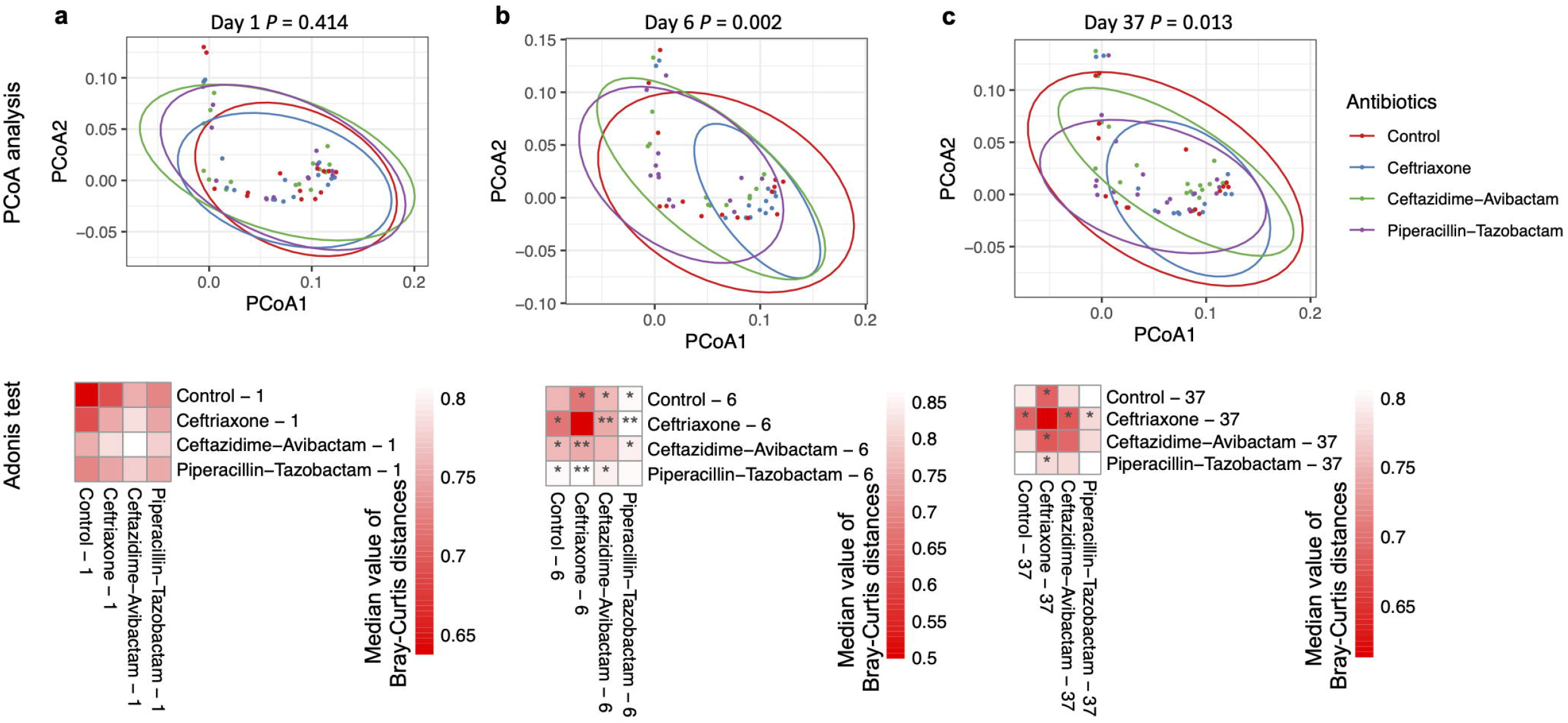
Effects of ceftriaxone, ceftazidime-avibactam, and piperacillin-tazobactam on the beta diversity of the gut microbiota. The impact of the three antibiotics on the gut microbiota was visualized using PCoA plots at days 1 **(a)**, 6 **(b)**, and 37 **(c)**. The significance of differences among the four groups was assessed using the Adonis test with *P*-values illustrated on the top of the PCoA plot. Pairwise comparisons were performed to determine statistical differences with *P*-values shown below the PCoA plot. * *P*-value ≤ 0.05, ** *P*-value ≤ 0.01, and *** *P*-value ≤ 0.001.

By day 37, the overall beta diversity differences among the four groups were less pronounced (P = 0.013) compared to day 6, yet remained significant (Figure 1c). Pairwise comparisons showed no significant differences between the ceftazidime-avibactam or piperacillin-tazobactam groups and the control group. However, the ceftriaxone group continued to exhibit significant differences from the other three groups (P ≤ 0.05), indicating that the effects of ceftriaxone on beta diversity persisted longer than those of the other two antibiotics.

### Effects of ceftriaxone, ceftazidime-avibactam, and piperacillin-tazobactam on composition of the gut microbiota

Using the LefSe test [21], differences in bacterial composition between antibiotic-treated groups and the control group at corresponding time points were assessed. The LDA scores indicated the levels of enrichment of specific bacterial taxa in a particular group compared to the corresponding control. The significance of the differences is presented in Figure S1. Clustering analysis, based on LDA scores of bacterial taxa exhibiting significant differences relative to controls, revealed intergroup similarities in the compositional changes of the gut microbiota across different time points and antibiotic treatments.

At baseline (day 1), minimal differences in bacterial composition were observed among the three antibiotic treatment groups (Figure 2). Consistent with the beta diversity analysis, compositional changes were most pronounced on day 6. The number of bacterial taxa exhibiting significant changes on day 37 exceeded that observed on day 1 but was lower than on day 6. Clustering analysis demonstrated that the effects of ceftazidime-avibactam and piperacillin-tazobactam on the composition of the gut microbiota, particularly on day 6, were more similar to each other than to those of ceftriaxone.

**Figure 2.**
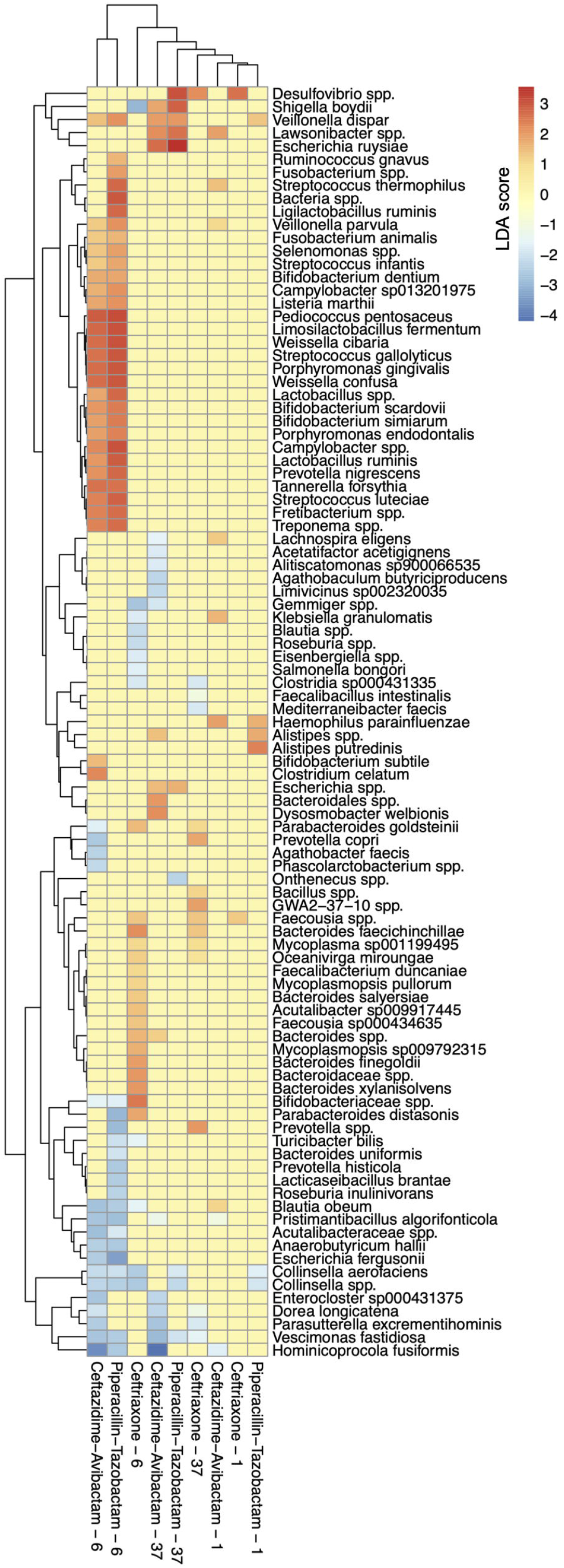
Effects of ceftriaxone, ceftazidime-avibactam, and piperacillin-tazobactam on the composition of the gut microbiota. The composition of the gut microbiota after treatment with the three antibiotics was compared to the corresponding control groups at the same time points using the LefSe analysis. The LDA score represents the degree of enrichment of specific bacterial taxa after antibiotic treatment relative to the control group. Clustering analysis was performed on all bacterial taxa that showed significantly different relative abundances compared to controls and were identified across the three time points and treatments, based on their LDA scores. The resulting heatmap illustrates the changes in gut microbiome composition over time and across antibiotic treatments, as well as the similarities between groups. *P*-values in the differential abundance analysis are detailed in Figure S1.

More specifically, a greater number of bacterial species were affected by ceftazidime-avibactam and piperacillin-tazobactam compared to ceftriaxone on day 6 (Figure S2 and Supplementary Dataset 2), indicating a broader inhibition spectrum for these two antibiotics [12]. Ceftriaxone predominantly promoted the enrichment of Gram-negative bacteria, whereas ceftazidime-avibactam and piperacillin-tazobactam enriched similar proportions of Gram-positive and Gram-negative bacterial taxa. These findings suggest that the impact of ceftazidime-avibactam and piperacillin-tazobactam on Gram-negative bacterial species differs from that of ceftriaxone.

### Effects of ceftriaxone, ceftazidime-avibactam, and piperacillin-tazobactam on bacterial interactions in the gut microbiota

Microbial homeostasis is governed not only by bacterial composition but by interactions among bacterial species. To evaluate these interactions, Spearman’s correlations between bacterial relative abundances were calculated, and significant correlations (FDR ≤ 0.05) were utilized to construct bacterial interaction networks (Figure S3). The degree, defined as the number of other bacteria interacting with a given bacterium, served as a measure of the strength of bacterial interactions.

On day 1, the degree distribution of bacteria in the ceftazidime-avibactam group differed significantly from that of the control group, while no significant differences were observed for the other two groups (Figure 3a). This suggests the presence of unknown baseline factors influencing bacterial interactions in the ceftazidime-avibactam group.

**Figure 3.**
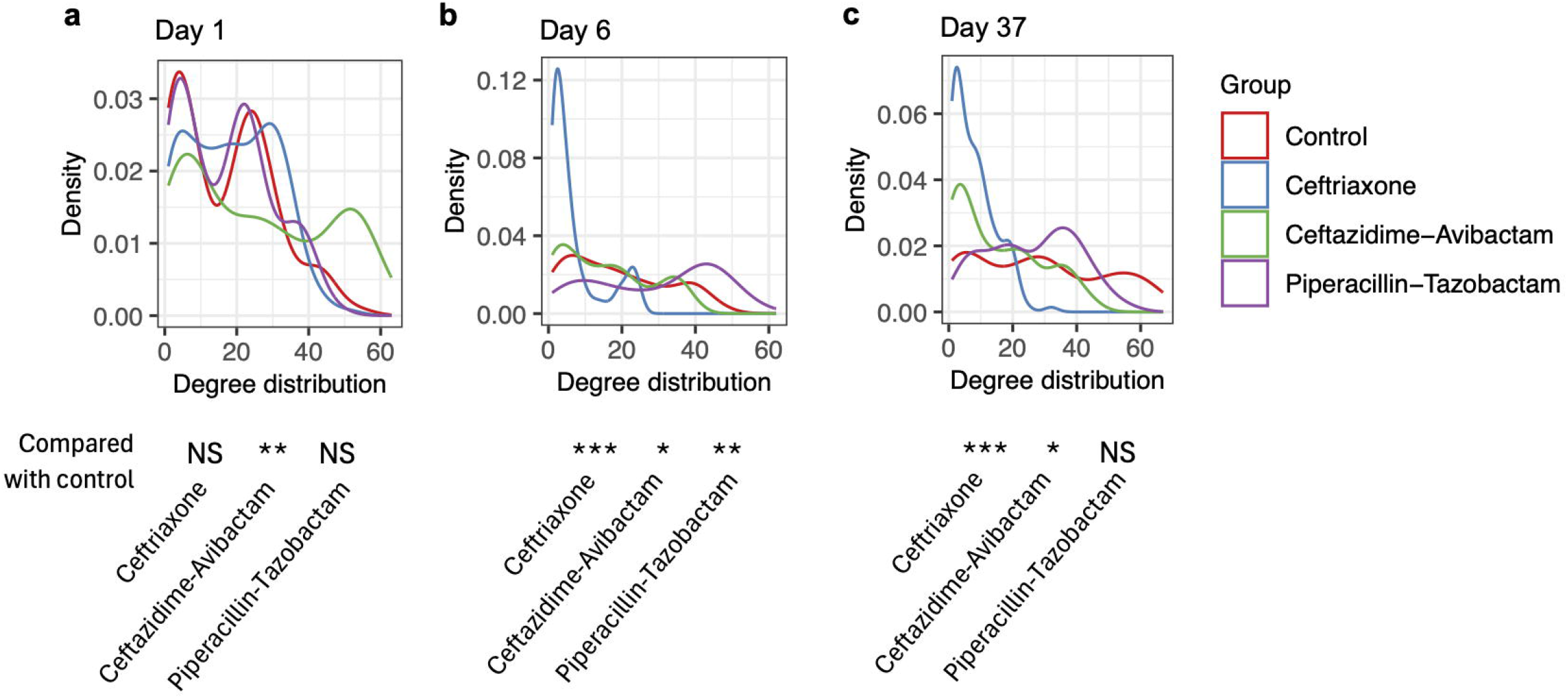
Effects of ceftriaxone, ceftazidime-avibactam, and piperacillin-tazobactam on the interactions among bacterial taxa in the gut microbiota. Spearman’s correlations between bacterial relative abundances were calculated, and significant correlations (FDR ≤ 0.05) were utilized to construct bacterial interaction networks (see Figure S3). The degree, defined as the number of other bacteria interacting with a given bacterium, served as a measure of the strength of bacterial interactions. The degree distributions of the gut microbiota on Day 1 **(a)**, 6 **(b)**, and 37 **(c)** are shown. The difference between each treatment and the corresponding control is tested using the two-sided Mann–Whitney U test. * *P*-value ≤ 0.05, ** *P*-value ≤ 0.01, and *** *P*-value ≤ 0.001.

Ceftriaxone had the most significant impact on bacterial interactions on day 6, which persisted through day 37 (Figures 3b and 3c). On day 6, the sizes of nodes, representing the degrees of bacterial taxa, for most bacterial taxa are smaller than those on day 1 (compare Ceftriaxone - 6 to Ceftriaxone - 1 on Figure S3). Therefore, most bacteria in the ceftriaxone group exhibited a degree of less than 8 on day 6, indicating disrupted interactions among bacteria (Figure 3b). This disruption remained significant on day 37 (P ≤ 0.001) (Figure 3c). However, the degree distribution of bacteria on day 37 was less affected than on day 6, as indicated by the reduced number of low-degree bacteria (compare values on the y-axis in Figure 3c versus Figure 3b).

On day 37, the ceftazidime-avibactam group still exhibited significant differences in bacterial interactions compared to the control group (P ≤ 0.05), although the differences were less pronounced than at baseline. No significant differences were observed between the piperacillin-tazobactam group and the control group, indicating that its effects on bacterial interactions were less persistent than those of the other antibiotics.

## Discussion

Based on the observed effects of ceftriaxone, ceftazidime-avibactam, and piperacillin-tazobactam on beta diversity, compositional changes, and bacterial interaction networks within the gut microbiota, it is evident that ceftriaxone exerts a stronger and more persistent influence on the gut microbiota compared to ceftazidime-avibactam and piperacillin-tazobactam. In contrast, ceftazidime-avibactam and piperacillin-tazobactam appear to have similar impacts on the gut microbiota. A similar phenomenon, in which ceftriaxone exerts a longer-lasting effect on the gut microbiota compared to piperacillin-tazobactam, has been observed in a murine model [22].

The stronger and more persistent influence of ceftriaxone on the gut microbiota could be attributed to several hypotheses. As mentioned earlier, ceftriaxone has a relatively longer half-life [6,9,10] of approximately 5.8 to 8.7 hours, maintaining prolonged pharmacological activity in the body. This slower clearance leads to extended intestinal residence time, increasing its sustained suppression of the gut microbiota, potentially resulting in more pronounced and sustained changes in the gut microbiota.

Another possibility is that the antibiotic spectra of ceftazidime-avibactam and piperacillin-tazobactam could be broader than that of ceftriaxone [11,12]. Particularly, ceftazidime-avibactam and piperacillin-tazobactam may exert a more balanced impact on Gram-positive and Gram-negative bacteria, as illustrated in Supplementary Figure S2. This balanced effect could promote the recovery of gut microbiota homeostasis. Additionally, other potential factors, such as differences in the metabolic capabilities of gut microbes to process these β-lactams, could also contribute to the observed differences in the impact of the three β-lactams on the gut microbiota.

Nevertheless, it is clear that dysbiosis of the gut microbiota is more severe following treatment with ceftriaxone. Thus, the risks of potential adverse health conditions associated with an imbalanced gut microbiota may be higher and persist for a longer duration after ceftriaxone administration compared to ceftazidime-avibactam and piperacillin-tazobactam. These findings could be beneficial for selecting appropriate antibiotics in clinical practice to minimize ecological disturbances in the gut microbiota. Alternatively, probiotics could be administered to patients receiving antibiotics that cause higher levels of disturbance to the human microbiota.

## Ethics statement

The study received approval from the Ethics Committee of the Third People’s Hospital of Cangnan County (No. II20250001). The publicly accessible dataset was sourced from the NCBI database (refer to Data availability for details), and information about informed consent to participate is available in the original publication linked to these dataset [20].

## Acknowledgments

We extend our gratitude to Dr. Bertrand Routy and his team at the Centre de Recherche du Centre Hospitalier de l’Université de Montréal, Canada, for generously sharing their public data.

## Author contributions

Guohang Huang conceived and designed the study. Guohang Huang and Ruyi Zhu performed the analyses. Guohang Huang wrote the paper. Lili Cai, Jinlin Guo, Ruyi Zhu, and Lina Zhu reviewed and edited the manuscript.

## Data availability

Publicly available datasets were downloaded from the NCBI SRA database (https://www.ncbi.nlm.nih.gov/sra) under the BioProject “PRJNA922086”.

## Ethics approval

The study received approval from the Ethics Committee of the Third People’s Hospital of Cangnan County (No. II20250001).

## Consent to participate

The publicly accessible dataset was sourced from the NCBI database (refer to Data availability for details), and information about informed consent to participate is available in the original publication linked to these dataset [20].

