## Supplementary Materials for "The influence of ceftriaxone, ceftazidime-avibactam, and piperacillin-tazobactam on the gut microbiota"

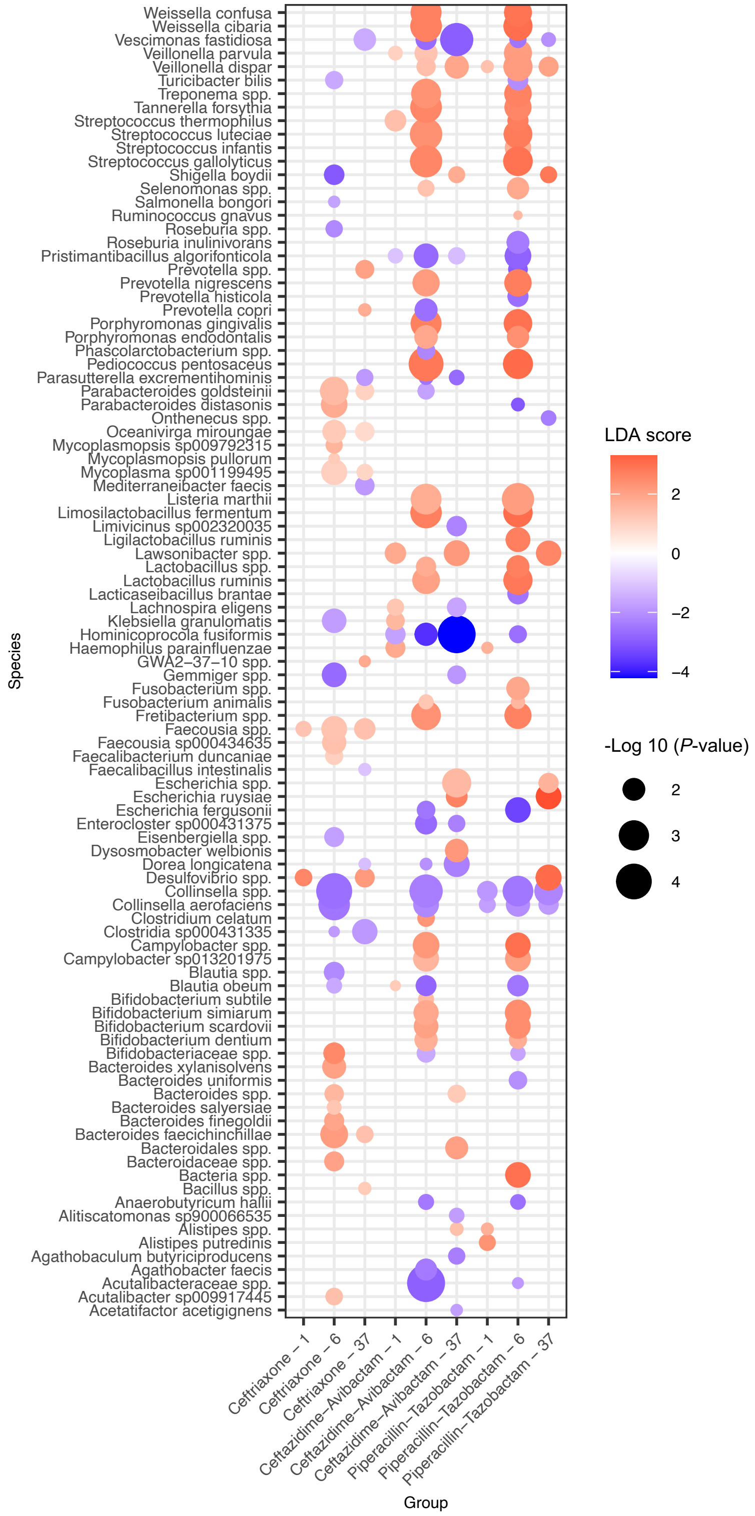


**Figure S1. Effects of ceftriaxone, ceftazidime-avibactam, and piperacillin-tazobactam on the composition of the gut microbiota.** The composition of the gut microbiota after treatment with the three antibiotics was compared to the corresponding control groups at the same time points using the LefSe analysis. The LDA score represents the degree of enrichment of specific bacterial taxa after antibiotic treatment relative to the control group.

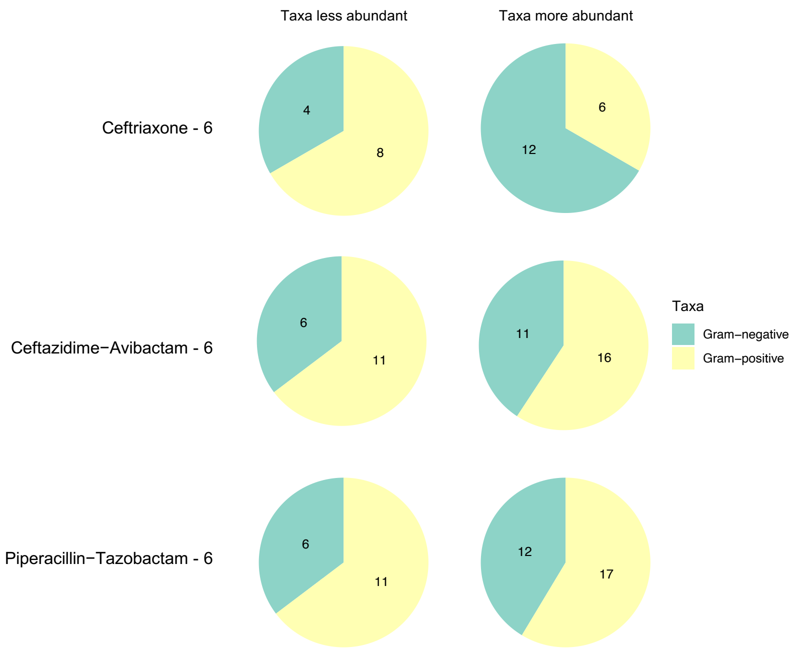


**Figure S2. The number of Gram-negative and Gram-positive bacterial taxa affected by treatments with ceftriaxone, ceftazidime-avibactam, and piperacillin-tazobactam.**

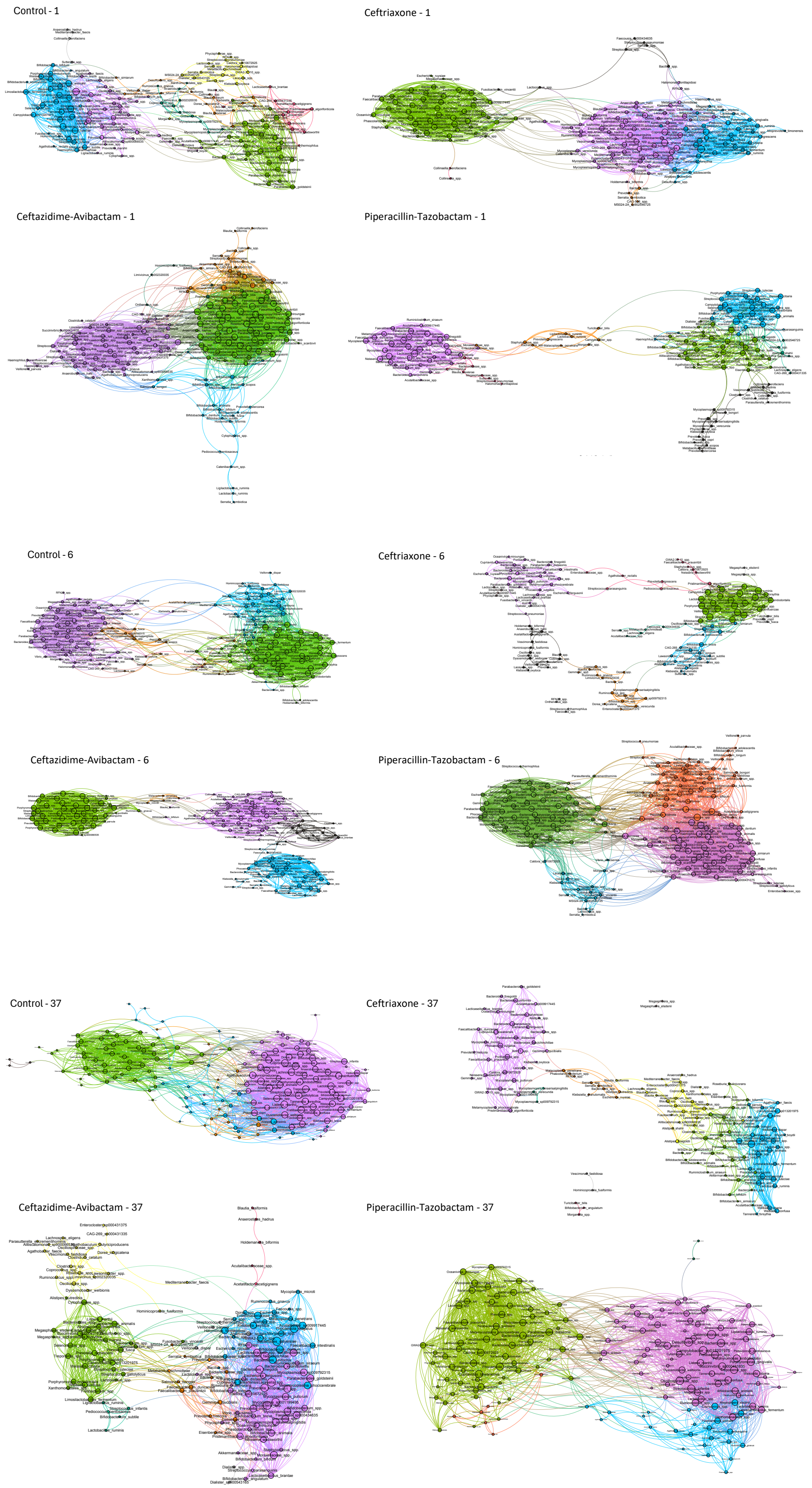
Control - 1
Ceftriaxone - 1
Ceftazidime-Avibactam - 1
Piperacillin-Tazobactam - 1
Control - 6
Ceftriaxone - 6
Ceftazidime-Avibactam - 6
Piperacillin-Tazobactam - 6
Control - 37
Ceftriaxone - 37
Ceftazidime-Avibactam - 37
Piperacillin-Tazobactam - 37

12

**Figure S3. Effects of ceftriaxone, ceftazidime-avibactam, and piperacillin-tazobactam on the interactions among bacterial taxa in the gut microbiota.** Spearman's correlation was used to evaluate associations between taxa pairs. The resulting *P*-values were corrected for multiple comparisons using the Benjamini-Hochberg method. Significant correlations (FDR ≤ 0.05) were used to construct unweighted bacterial interaction networks in Gephi. The degree of each bacterial node was quantified using the 'Avg. Path Length' function and determined the size of each node, while modules within the networks were identified using the 'Modularity' function. Nodes were color-coded according to their assigned module.
